# A Neural Circuit Model of Proprioceptive Feedback from Muscles Recruited during an Isometric Knee Extension

**DOI:** 10.1101/802736

**Authors:** Gareth York, Hugh Osborne, Piyanee Sriya, Sarah Astill, Marc de Kamps, Samit Chakrabarty

## Abstract

The influence of proprioceptive feedback on muscle activity during isometric tasks is the subject of conflicting studies. To better understand the relationship, we performed an isometric knee extension task experiment at four pre-set angles of the knee, recording from five muscles, and for two different hip positions. We applied muscle synergy analysis using NMF on the sEMG recordings to identify structure in the data which changed with internal knee angle, suggesting a link between proprioception and muscle activity. We hypothesised that such patterns in the data arise from the way proprioceptive and cortical signals are integrated in neural circuits of the spinal cord. Using the MIIND neural simulation platform, we developed a computational model based on current understanding of spinal circuits with an adjustable proprioceptive input. The model produces the same synergy patterns as observed in the experimental data indicating that such synergies are indeed encoded in the neural connectivity of the spinal cord and modulated by the proprioceptive input. When matching the proprioceptive input to the knee angles of the experiment, the model predicts the need for three distinct inputs: two to control the normal reciprocity between the agonist and antagonist muscles, and an additional to match the non-linear trend towards the limit of extension. Finally, we discuss the risks of using NMF for synergy analysis and demonstrate how to increase confidence in its results. Future modelling of human motor outputs should include interneuronal spinal circuits such as this to distinguish the modulatory role of supraspinal and peripheral afferent inputs to the spinal cord, during both passive and dynamic tasks.

**Significance statement:** Sensory feedback from muscles has a significant role in motor control, but its role in tasks where limbs are held in a fixed position is disputed, because the effect is reduced when muscles are not stretched. Here, we first identified patterns of muscle activity during such tasks which changed with different leg positions. We developed a computational spinal motor circuit model with adjustable muscle stretch input, which reproduced the same patterns of activity as observed experimentally. The model predicts three distinct muscle stretch signals required to produce the activity patterns for all leg positions. Because the connections in the model are based on well known spinal circuits, it is likely the observed activity patterns are generated in the spinal cord.

## 1. Introduction

The execution of a motor task is achieved through the integration of simple movement commands which are modulated by sensory feedback from the periphery over time. The role of proprioceptive feedback in the recruitment of muscle fibres to counter load during a given task is well understood. However, its role in control of muscle activity especially in the commonly tested static task involving a single joint, is still poorly understood. The benefit of investigating control in an isometric task, where the limb is supported in place, is that the influence of proprioceptive signals related to limb movement, stretch activated Ia afferent for example, can be eliminated. There is evidence that other proprioceptive afferents like the Golgi-tendon organ (Ib) may still contribute during an isometric task. For example, neuromechanical modelling (Kistemaker et al. 2013) has been used to demonstrate the importance of Ib afferent signals in motor control. Furthermore, in the study of a de-afferented man (Rothwell et al. 1982), isometric control was shown to be impaired in some tasks. However, a previous report on activation patterns in muscles of the upper arm during an isometric task (Roh, Rymer, and Beer 2012) showed no change when the arm position was altered. A study of muscles in the cat hind limb during a balance task (Torres-Oviedo, Macpherson, and Ting 2006) showed an in-variance to starting position.

A common technique for identifying the effect of proprioception on motor execution is the use of muscle synergy analysis. There is great variation in the way a motor task can be performed, even at a single joint. Each variation is produced from a combination of muscle recruitment patterns, often described in terms of time. These patterns are commonly referred to as muscle synergies. Their use by the central nervous system (CNS) to alleviate the degrees of freedom problem is accepted but little is known about the mechanism of their recruitment (Grillner 1985; Bizzi, Mussa-Ivaldi, and S. Giszter 1991; Tresch, Saltiel, et al. 2002). Similar synergies are reported across species, especially for routine repetitive tasks like locomotion in vertebrates (Yang, Logan, and S. F. Giszter 2019). As well as providing insight directly, synergy patterns give a clear summary of the structure of experimental data (usually electromyographic recordings) and are therefore a good choice for identifying changes and trends due to differing conditions.

In this study, we perform an isometric knee extension task experiment in which the leg is held extended in a brace at a set angle, minimising activity across all joints in the subject. To isolate the modulation of the spinal motor execution circuits, subjects voluntarily maximised activity of the rectus femoris muscle during recordings thereby limiting the influence of the descending drive. For each trial, the brace was set at a distinct internal knee angle so the length of the muscles were different before the recordings began. Activity of the muscles at the knee were recorded, as surface electromyograms (sEMG) and synergies during the task were computed for all angles with two different hip positions. Our primary finding is that the activity pattern changes with the internal knee angle, suggesting the altered muscle length (and thereby the difference in proprioceptive input) does play a role. The specific patterns and trends observed in the synergies were then reproduced using a neuronal model, to ascertain the role of proprioceptive inputs in recruitment of the synergies. The synergies were identified using non-negative matrix factorization (NMF), which has previously been used in other muscle synergy studies (Roh, Rymer, and Beer 2012; Torres-Oviedo, Macpherson, and Ting 2006; Saltiel et al. 2001) although recently, aspects of the technique have come under some scrutiny (Barradas et al. 2020). In addition to demonstrating a model which successfully reproduces the synergies, we also discuss two features in our NMF results which gives us greater confidence in our conclusions.

The model is described at the level of populations of neurons and is simulated using population density techniques (PDTs) which have been shown to accurately model population aggregates (like firing rates) while retaining a close correspondence to spiking neurons -more than so called neural mass models -without producing the overhead of simulating thousands of neurons. MIIND (De Kamps, Lepperød, and Lai 2019) is a simulation platform which uses a PDT and was chosen for its support for fast prototyping of population circuits as well as its compatibility with the Python programming language with which all data analysis was performed.

Building the neural model gives a number of advantages including a mechanistic explanation of the experimental results over and above a simple observation. It also provides a prediction of the existence of multiple proprioceptive signals combining to produce the observed behaviour at different knee angles. Finally, the model is based on well understood spinal circuits (Pratt and Jordan 1987; Jankowska et al. 1967) and can therefore be easily integrated into other models which require a proprioceptive input at the spinal cord level. Examples of such models are described in the discussion section.

## 2. Materials and Methods

### 2.1 Ethics

The study was conducted according to the Declaration of Helsinki and all experimental protocols were approved by the University of Leeds Research Ethics Committee (reference number BIOSCI 16-004). Healthy subjects (n= 17, male = 9, female = 8) with an age range of 18-30 (24.4± 2.57 years) were recruited to participate in this study. Exclusion criteria included previous knee or leg injuries, if participants had done exercise within 48 hours prior to testing, knee stiffness or self-reported pain, use of recreational or performance enhancing drugs, ingested alcohol in the previous 24 hours or were unable to provide informed consent. Subjects provided informed written consent to the study, noting possible risks associated with the activity.

### 2.2 Data Collection

Surface sEMG was recorded from seven muscles of the subjects dominant leg; rectus femoris (RF), vastus lateralis (VL), vastus medialis (VM), semitendinosus (ST), biceps femoris (BF), medial gastrocnemius (MG) and tibialis anterior (TA) – of which the MG and TA were discarded from further analysis, due to low signal to noise ratio. Data analysis was therefore performed on the five remaining muscle recordings. The skin was prepared for electrodes with shaving, cleaning with alcohol wipes and then application of conductive electrode gel. Data was sampled at 2 kHz using wireless Delsys Trigno IM electrodes. Electrodes were placed on the muscle belly, defined by landmarks as described in Rainoldi, Melchiorri, and Caruso 2004 based on anatomical observations: VL-between the greater trochanter and the lateral epicondyle; VM -on the distal fifth of the medial knee joint; RF -between the greater trochanter and the lateral epicondyle; VM -on the distal fifth of the medial knee joint; RF -between the anterior superior iliac spine and the superior pole of the patellar, and MG belly located in distal third of the medial knee joint.

### 2.3 Experimental Protocol

Subjects were asked to lay on a standard medical examination bed. They were then shown how to perform an isometric knee extension with the leg brace supporting their dominant leg. Subjects were provided the resulting sEMG output recorded from RF as a feedback to help them learn the performance of the task. Subjects were asked to perform an isometric knee extension at maximal voluntary effort for five seconds, attempting to maximise RF activity. They were shown the RF sEMG output to aid them. This was repeated six times with a three minute rest between contractions. The dominant knee was fixed at one of four angles using a Donjoy TROM locking knee brace at 0°, 20°, 60° and 90°. The angle of the knee was always measured against the hip joint and the bony prominence on the outside of the ankle. Data was collected in two different positions and sessions for each subject. In position one, the participant was supine with both legs flat against the bed. In position two the contralateral leg was kept bent such that the foot is flat against the bed so that both the knee and hip are fully flexed. The position selected for each subject was randomized for their first session. In the second session, the subject performed the task in the other position.

### 2.4 Data Preprocessing

Data was prepared for NMF via a three stage filtering process. Signals were initially band-pass filtered (high pass = 20 Hz, low pass = 450 Hz, second order Butterworth filter), rectified and then finally zero-lag high-pass filtered (5 Hz, second order Butterworth filter) to remove frequency changes induced by rectification. Finally, a simple moving average was applied with a time window of 0.5s to further smooth the signals. Each sEMG channel was normalized to the maximum value for that channel across all six contractions. Data was visually inspected and segmented into equal sections containing one burst. NMF was performed on all contractions across all sEMG channels.

### 2.5 Synergy Extraction

We used NMF to identify muscle synergies during the task. Information theory shows that this dimensionality reduction reflects latent structure in the data, which we interpret as muscle synergies (Lee and Seung 2001). NMF’s chief advantage compared to other approaches is the constraint of non-negativity aligning with muscle activity i.e. muscle activation is never negative. NMF is also more effective at identifying latent structure in the data when compared to other techniques such as principal component analysis (Ebied et al. 2018).

The five raw (but smoothed) sEMG time series were combined into a matrix *D* of size 5*× n* where *n* is the length of the time series. We used iterative NMF decomposition algorithms (Tresch and Bizzi 1999; Lee and Seung 2001) to reduce D to a combination of two matrices, *W* and *C* such that,

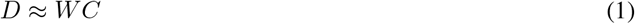

C is an *N× n* matrix where *N* is the chosen NMF rank factor, in this case, 2. Each row of *C* represents some structure in the time series similar to a PCA component. *W* is a 5*× N* matrix which, when multiplied by *C*, approximates matrix *D*. Each column of *W* quantifies the amount that the corresponding row in *C* contributes to the original data in *D* (Lee and Seung 2001; Donoho and Stodden 2004; Berry et al. 2007; Torres-Oviedo and Ting 2007). Each synergy, *s*, is represented by the corresponding column *W∗*_*s*_ and row *C*_*s*_*∗*. We describe *C*_*s*_ as the activation pattern of the synergy as it represents some underlying structure of the original sEMG time series. We refer to *W ∗*_*s*_ as the muscle contribution vector of the synergy as each component magnitude indicates the contribution of the synergy’s activation pattern to the associated muscle activity.

Selection of rank factor is critical to achieving dimensionality reduction such that *C* has fewer rows than *D*. Rank factor was chosen consistent with previous literature (Tresch, Cheung, and d’Avella 2006) such that rank factor was increased to the minimum required for the variance accounted for (VAF) by *WC* compared to *D* was greater than 90%. VAF was calculated for each synergy profile for both the individual muscle and for all muscles collectively. If VAF was below 90%, the resulting synergies were discarded. The iterative optimization algorithm used was initialized using singular value decomposition to reduce calculation time and to ensure a unique and reproducible result (Boutsidis and Gallopoulos 2008). Each row in *C* and column in *W* was normalized to its maximum value. Cosine similarity analysis was used in a pairwise fashion to determine the similarity between subjects’ synergy vectors and activation coefficients *W*_*∗s*_ and *C*_*s∗*_ (Rimini, Agostini, and Knaflitz 2017).

### 2.6 Neural Population Modelling

We aimed to create a neural population model such that applying NMF to the firing rate activity of the motor neuron populations would yield the same synergy patterns as those identified in the sEMG data. We did not attempt to reproduce simulated sEMG signals. Instead, we assumed that the cumulative activity of multiple motor units described by the average activity of distinct motor neuron populations would serve as a proxy for sEMG. When designing the model, we considered rate-based models which represent a population metric, for example the average firing rate (Wilson and Cowan 1972) or oscillation frequency (Kuramoto 1991), abstracted from the underlying individual neurons. Rate-based models are suitable for reproducing firing rates in neural circuits, but there is no clear relationship with the state of the underlying neural substrate. Although not essential for this study, in light of more detailed spinal models used in the field where individual neurons are simulated (McCrea and Rybak 2008), as well as future development of the modelling work, we are interested in a technique that retains a closer relationship with the state of the spiking neurons that comprise the neural circuit. PDTs are such a technique: they retain information about the state of neurons in the circuits but calculate population level aggregates directly.

### 2.7 Population Density Techniques

Population Density Techniques (PDTs) model neural circuits in terms of homogeneous populations of neurons. The individual neurons are described by a model, in this case, exponential integrate-and-fire. The model of an individual neuron is characterised by a so-called state space: the values that determine the state of individual spiking neurons. For a simple neuron model, this can be its membrane potential. For more complex models, variables such as the state of a synapse can be included. PDTs represent a population by a single density function that indicates how neurons are distributed across the neuron model’s state space.

### 2.8 MIIND

MIIND is a neural simulator (De Kamps, Baier, et al. 2008) which implements a version of a PDT to simulate multiple interacting populations of neurons. It can provide a visual representation of the probability density function by displaying the density during simulation. Figure 5A shows an example of this visual representation.

A network of populations can be built in MIIND using a simple XML style code format to list the individual populations and the connections between them. Populations in the network interact via their average firing rates, which are assumed to be Poisson distributed spike trains. For each connection, the firing rate of the source population becomes the average rate of the Poisson distributed input spikes to the destination population. The connections defined in the XML code have three parameters: the post-synaptic potential or instantaneous synaptic efficacy, the number of individual connections between source neurons and target neurons, and a delay which can be used to approximate time taken for spike propagation and synapse transmission.

### 2.9 The Spinal Circuit Model

We used MIIND to build a network of populations of exponential integrate and fire (EIF) neurons according to the connectivity diagram in Figure 1. Table 1 shows the connection parameters for all populations in the model. All populations use the same underlying neuron model as described in Equation 2.

**Figure 1:**
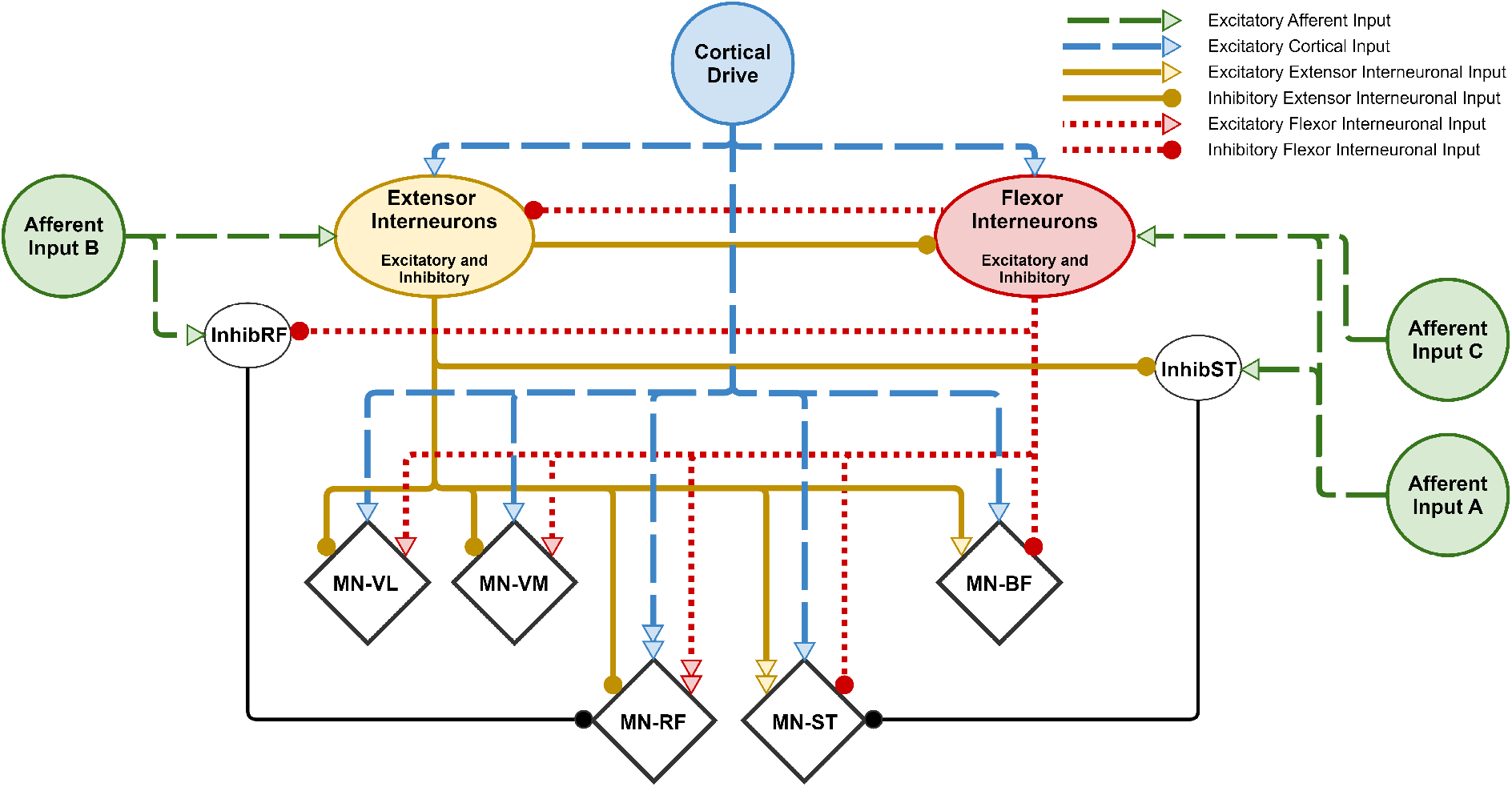
Schematic of connections between simulated spinal populations. MN-VL (Vastus Lateralis), MN-VM (Vastus Medialis), MN-RF (Rectus Femoris), MN-ST (Semitendinosus) and MN-BF (Biceps Femoris) motor neuron populations are identified as diamonds although all populations consist of EIF neurons. The Extensor and Flexor Interneuron populations allow both outgoing excitatory and inhibitory connections. All populations receive a background level of input producing a baseline activity. Parameters for the network connectivity are provided in Table 1. MN-RF and MN-ST receive much higher excitation from the interneuron populations than the other motor neuron populations (indicated as a double arrow). The InhibRF and InhibST populations are used to offset the level of bias given to the MN-RF and MN-ST populations. Afferent Input A and B control the balance of input to the Flexor and Extensor Interneuron populations respectively which influences the agonist/antagonist bias in synergy 2. Afferent Input C represents an additional input, activated at the limit of extension, further increasing the excitatory input (via the flexor interneurons) to the agonist motor neuron populations. Connections that exist in other models but which are not required to produce the observed synergies have been omitted. For example, direct afferent inputs to motor neuron populations. The relative strengths of each connection are also not shown (except for MN-RF and MN-ST) but can be found in the connectivity parameters (Table 1).

**Table 1:** Parameters relevant to each connection between populations and from inputs in the model. Values for input activity are provided in the form of an average firing rate.

| Population name | Source population name | Post synaptic delta efficacy (mV) | Average number of incoming connections to each neuron | Connection delay time (ms) | Average firing rate where defined (Hz) |
| --- | --- | --- | --- | --- | --- |
| MN-RF | Extensor Interneurons | -0.052 | 50 | 2 |  |
| MN-VL | Extensor Interneurons | -0.052 | 50 | 2 |  |
| MN-VM | Extensor Interneurons | -0.052 | 50 | 2 |  |
| MN-ST | Extensor Interneurons | 0.052 | 100 | 2 |  |
| MN-BF | Extensor Interneurons | 0.052 | 50 | 2 |  |
| MN-RF | Flexor Interneurons | 0.052 | 100 | 2 |  |
| MN-VL | Flexor Interneurons | 0.052 | 50 | 2 |  |
| MN-VM | Flexor Interneurons | 0.052 | 50 | 2 |  |
| MN-ST | Flexor Interneurons | -0.052 | 50 | 2 |  |
| MN-BF | Flexor Interneurons | -0.052 | 50 | 2 |  |
| MN-RF | InhibRF | -0.052 | 50 | 2 |  |
| MN-ST | InhibST | -0.052 | 50 | 2 |  |
| InhibST | Extensor Interneurons | -0.052 | 50 | 2 |  |
| InhibRF | Flexor Interneurons | -0.052 | 50 | 2 |  |
| Extensor Interneurons | Flexor Interneurons | -0.052 | 50 | 2 |  |
| Flexor Interneurons | Extensor Interneurons | -0.052 | 50 | 2 |  |
| Extensor Interneurons | Background | 0.1 | 100 | 0 | 450 |
| Flexor Interneurons | Background | 0.1 | 100 | 0 | 450 |
| InhibST | Background | 0.1 | 100 | 0 | 450 |
| InhibRF | Background | 0.1 | 100 | 0 | 350 |
| MN-RF | Background | 0.1 | 100 | 0 | 350 |
| MN-VL | Background | 0.1 | 100 | 0 | 350 |
| MN-VM | Background | 0.1 | 100 | 0 | 350 |
| MN-ST | Background | 0.1 | 100 | 0 | 350 |
| MN-BF | Background | 0.1 | 100 | 0 | 350 |
| MN-RF | Cortical Drive | 0.1 | 100 | 0 | 60* |
| MN-VL | Cortical Drive | 0.1 | 100 | 0 | 50 |
| MN-VM | Cortical Drive | 0.1 | 100 | 0 | 50 |
| MN-ST | Cortical Drive | 0.1 | 100 | 0 | 50 |
| MN-BF | Cortical Drive | 0.1 | 100 | 0 | 50 |
| Extensor Interneurons | Cortical Drive | 0.1 | 100 | 0 | 50 |
| Flexor Interneurons | Cortical Drive | 0.1 | 100 | 0 | 50 |
| Extensor Interneurons | Afferent Input B | 0.1 | 100 | 0 | 50 |
| InhibRF | Afferent Input B | 0.1 | 100 | 0 | 0-100** |
| Flexor Interneurons | Afferent Input A | 0.1 | 100 | 0 | 100-300 |
| InhibST | Afferent Input A | 0.1 | 100 | 0 | 0-200 |
| Flexor Interneurons | Afferent Input C | 0.1 | 100 | 0 | 0 or 100 |

\* During the task, cortical drive input transitions from 0Hz to these values then back to 0Hz
\*\* During the task, afferent activity transitions from 0Hz to values in this range depending on the level of afferent feedback

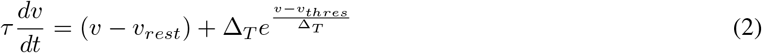

Where *v* is the membrane potential, *v*_*rest*_ = − 70 mV, Δ_*T*_ = 1.48, *v*_*thres*_ = *—*56 mV, and 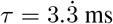. The parameters were chosen so that populations could produce a wide range of average firing rates between 0 and 200 Hz to exhibit typical neuronal frequencies. We chose to use an EIF model in contrast to more commonly used Hodgkin Huxley style neurons. This is because the objective was not to reproduce the sEMG signals exactly, but to provide a concise explanation for the overall synergy patterns. We expected that any particular description of activation of ion channels (as in a Hodgkin Huxley style model) would have no significant impact on the population level activity or synergy patterns in this task and would therefore dilute the power of the model. The main structure of the network consists of two neural populations, named “Extensor Interneurons” and “Flexor Interneurons”, connected together in a network with five motor neuron populations named MN-RF, MN-VL, MV-VM, MN-ST, and MN-BF for each respective muscle. The extensor and flexor interneuron populations represent combinations of excitatory and inhibitory neurons and therefore can project both kinds of connections to other populations in the network. Other studies have previously described the connection motif of agonist inhibition with antagonist excitation. This is used here to connect the interneuron and motor neuron populations to elicit the agonist/antagonist relationship between the five muscles (**bigland1981sEMG**; Sherrington 1909; Doss and Karpovich 1965; Pierrot-Deseilligny and Burke 2005). These features, including the mutual inhibition between the two interneuron populations, also appear in the central pattern generator (CPG) model of McCrea and Rybak 2008. Although the rhythm and pattern formation parts are not included, the implications for applying a CPG model to an isometric task are discussed later.

#### 2.9.1 Cortical Drive

All supraspinal activity comes from the cortical drive input and is responsible for the “contraction”. There is a direct connection to all motor neuron populations but a higher level of drive is provided to the MN-RF motor neuron population indicative of the muscle which is being maximally contracted in the task. Cortical drive also projects to the extensor and flexor interneuron populations. During the simulation, the input to the two interneuron populations and motor neuron populations begins at 0Hz before increasing to 50Hz (MN-RF to 60Hz) over 1 second, then five seconds later, dropping back to 0Hz over 1 second. The average firing rate of each of the five motor neuron populations was recorded at a rate of 10 kHz (corresponding to the 0.1ms time step of the simulation) then sampled at 2ms intervals. NMF was performed on the resultant time series as described for the experimental recordings.

#### 2.9.2 Afferent Inputs to the Network

We hypothesised that the most important factor in shaping the observed synergies would be the connectivity of the network model and that they could be modulated with a proprioceptive input. As demonstrated by the variation in synergy patterns of the sEMG recordings, there are potentially many functionally separate afferent signals contributing to muscle activity at different angles and positions. To capture the most identifiable patterns (the agonist/antagonist bias with changing knee angle and near maximum extension), we introduced three different afferent inputs to the population network. Afferent inputs A and B allow us to change the balance of activity between the extensor interneuron and flexor interneuron populations. Afferent input C provides additional input to the flexor interneuron population to represent a functionally different signal when near maximum extension.

#### 2.9.3 Synergy Bias for the Bifunctional Muscles

Finally, we augmented the network to control the bias in many of the synergy patterns towards the two bifunctional muscles, RF and ST. The strength of the connections from the extensor interneuron population to MN-ST and flexor interneuron population to MN-RF was increased, effectively increasing the number of connections between those populations. To modulate the effect of the afferent feedback input on these connections and to reproduce the change in the bias with change in knee angle, two additional populations of inhibitory neurons were added to the model: InhibST and InhibRF. This network motif of an additional excitatory drive coupled with a controllable inhibitory input has previously been used to reproduce observed activity in semitendinosus and rectus femoris of a cat (Shevtsova et al. 2016). We believe this to be the first time such a circuit has been applied to human studies.

### 2.10 Statistical Analysis

All statistical analysis was performed in Python 3.6.2. Cosine similarity analysis was used to compare sEMG profiles, synergy activation patterns and muscle contribution vectors. Cosine similarity analysis is sensitive to differences in vectors that may have equal variation.

### 2.11 Code Accessibility

NMF analysis and cosine similarity analysis was performed using a custom designed program in Python 3.6.2. MIIND is available at [URL redacted for double-blind review] and the model files and simulation results are accessible at [URL redacted for double-blind review].

## 3 Results

We recorded from 7 different muscles of the leg, but only 5 of these were used for further analysis of activity patterns as on examination the muscles TA and MG were always inactive, as expected due to the nature of the task. We extracted muscle synergies from the sEMGs recorded from the 5 different muscles across the two different positions to examine how proprioceptive feedback alters muscle synergy recruitment. The contralateral hip was flexed or relaxed to induce passive insufficiency in the recorded leg to highlight differences in proprioceptive feedback. A spinal population circuit model was created with connections between interneurons, motor neurons and afferent feedback, based on current CPG models and accepted neural circuits (Pierrot-Deseilligny and Burke 2005). Similarities between the synergies observed experimentally and those extracted from the model were compared. Figure 5B shows the sEMG signals for all five muscles after rectification and smoothing, and the average firing rates of the motor neuron populations in the simulation. It is difficult to discern, by eye, the differences between the time series, highlighting the need for analysis techniques such as NMF.

### 3.1 NMF identifies two muscle synergies from the sEMG activity

To identify synergies appropriate for experiment-model comparison, NMF was performed with a range of rank values, one rank per synergy. The appropriate rank to use was chosen as the number required to raise the VAF above 90% (Figure 2). In this case, rank two raised VAF above this threshold. Although 90% is an arbitrary threshold, and there are other methods for choosing appropriate rank, patterns identified by three or more synergies were less consistent across participants. As described in section 2.4, each synergy consists of a column of matrix *W* with length five (one value per muscle) and a row of matrix *C* representing a time series describing some underlying structure of the original data. For each muscle, the corresponding component of *W* _*∗s*_ multiplied by *C*_*s∗*_, gives the contribution of synergy, *s*, to that muscle’s sEMG. Cosine similarity analysis was performed on the synergy rows and columns across participants for each position, synergy and angle. There is high correlation between synergy 1 results among the participants, regardless of position and internal knee angle (table in figure 2). Though not as high as synergy 1, there is also high correlation between participants for synergy 2. Despite some variation, there is a common pattern of muscle synergy recruitment across all participants.

**Figure 2:**
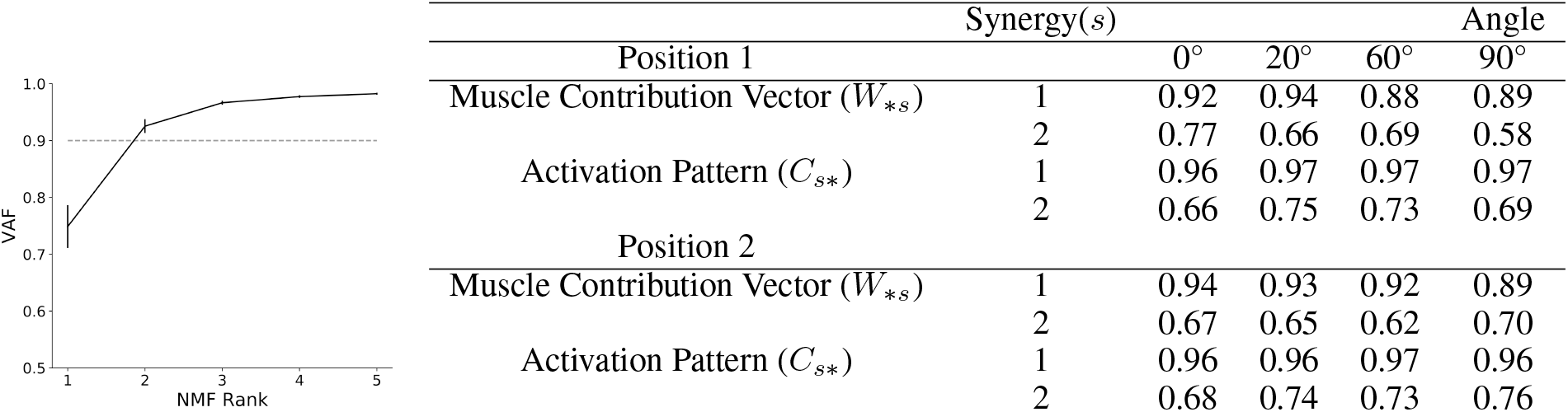
Average VAF scree plot for rank one to five NMF dimensionality reduction across all angles and both positions of the isometric knee extension task. The 90% VAF threshold indicates that two is the appropriate rank to use and therefore the number of synergies to extract. Error bars show Standard Error of the Mean (SEM). In the table, synergy rows (activation patterns) and columns (contribution vectors as defined in section 2.4) were compared across all pairs of participants using cosine similarity analysis giving a value between 0 (uncorrelated) and 1 (Highly correlated). For both positions (activating or inactivating the contralateral hip flexors) and for all internal knee angles, there is high correlation between subjects indicating that, during the task, the same synergy patterns are being recruited by the majority of subjects.

### 3.2 Proprioceptive feedback alters muscle synergy structure in a manner reflecting muscle stretch

#### 3.2.1 Synergy 1 : There is a pattern showing coordinated, balanced recruitment of all five muscles in synergy 1

The NMF process generates, for each of the two synergies, a time series activation pattern and a vector of five values, one for each muscle. Figure 3 shows the vector and time series of synergy 1 (A) and 2 (B) for both positions across different internal knee angles. The activation patterns (line plots) should be considered in conjunction with the five value muscle contribution vectors shown in the bar charts. Synergy 1 represents co-activation of all muscles as would be expected in an isometric extension and contributes to the majority of the observed sEMG activity. Because of this, the activation pattern closely matches the overall profile observed in the raw sEMG data (the transition from low to high to low activity during the contraction). The high muscle contribution values for all five muscles indicates that this activation pattern is present in all five sEMG recordings. Both the activation pattern and muscle contribution weights are well conserved across all angles, positions, and muscle groups. Among the muscle contribution values, there is a slight shift between 0° and 90° from a quadriceps bias to a hamstrings bias but this change is less apparent in position 2. Because the vector values are very well correlated across participants in synergy 1, any individual trial in which the synergy 1 vector values were not all at a high level (i.e. the associated contribution value of one or more muscles was unexpectedly low) could be eliminated as an outlier before analysis of synergy 2 was performed. Therefore, if any vector value in a trial fell outside the interquartile range of the other four values, the trial was eliminated.

**Figure 3:**
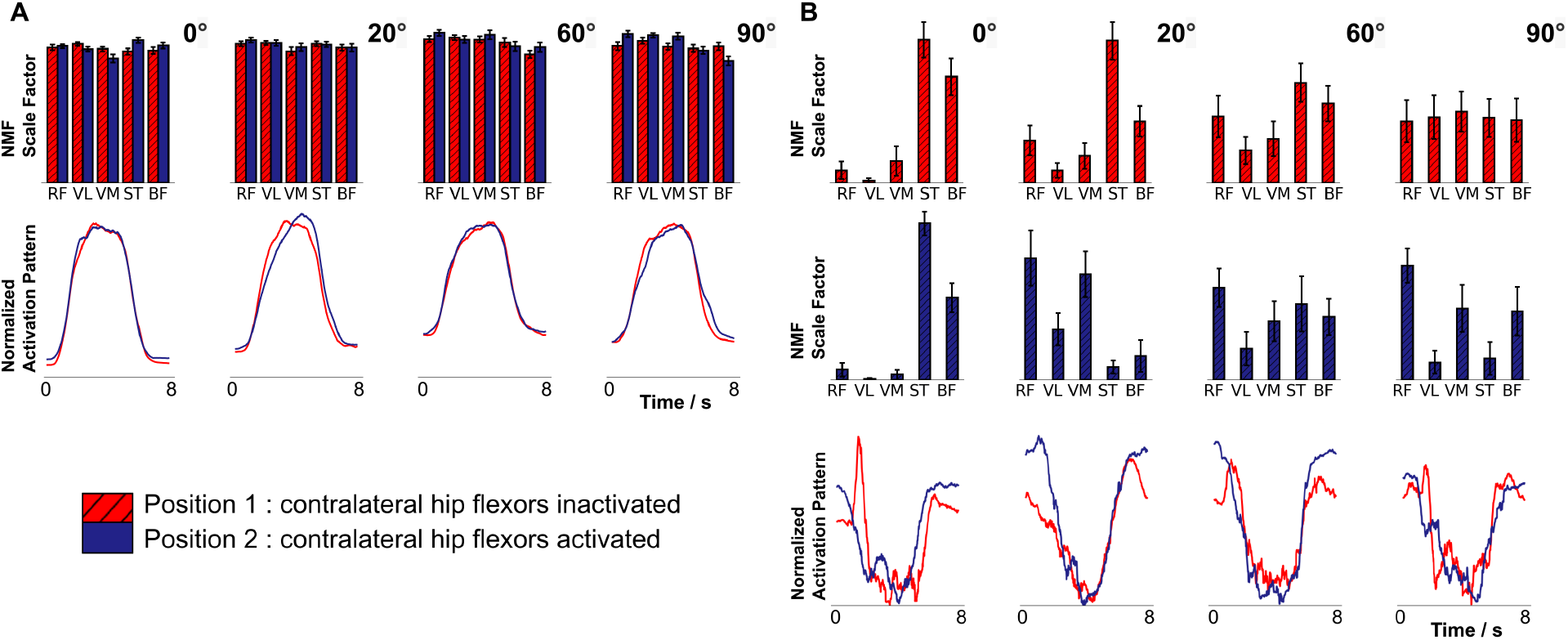
Muscle synergies extracted using rank two NMF from an isometric knee extension task at four internal angles of the knee (0°, 20°, 60°, and 90°) (N = 17, mixed gender, male = 9, female = 8, age range of 18-30 (24.4±2.57 years). Subjects performed 6 contractions of 5s with the subject being asked to maximize rectus femoris activity. NMF was performed on the average sEMG of each subject’s 6 contractions. The experiment was repeated across two positions inactivating (red values) or activating (blue) contralateral hip flexors. Line charts are activation patterns identified by NMF as underlying structure in the original sEMG time series. Bar charts show the contribution of the associated activation pattern to the activity of each of the five muscles in arbitrary units. Error bars represent standard error of the mean. A: Synergy 1 demonstrates a balanced, coordinated contraction across muscle groups in line with what is observed in the raw sEMG data. B: Synergy 2 for position 1 (top in red) shows the balancing of antagonist/agonist activations between the quadriceps muscles and hamstrings. Synergy 2 for position 2 (middle in blue) shows the same extreme antagonist bias at 0° with the leg at maximal extension. However, there is a strong agonist bias at 20° perhaps demonstrating the effect of less passive insufficiency. RF:Rectus Femoris; VL:Vastus Lateralis; VM:Vastus Medialis; ST:Semitendinosus; BF:Biceps Femoris.

#### 3.2.2 Synergy 2 : Changing the internal knee angle alters the contribution of the activation pattern in the agonist and antagonist muscles

The shape of the activation pattern of synergy 2 indicates that this synergy captures a difference between the activity of each muscle during the contraction and at rest. During the contraction, the contribution of synergy 2 reduces to near zero because all muscles reach their maximal activity which is the feature captured solely by synergy 1. The contribution vectors of synergy 2 change for different knee angles and position indicating that there is a proprioceptive effect on muscle activation even in an isometric task such as this.

In position 1, where the contralateral leg is straight, as the internal knee angle of the recorded leg is increased there is a marked drop in the average of contribution vector magnitudes for the antagonist muscles whereas the average of the agonist muscles shows the opposite trend. At 90°, magnitudes are similar for all muscles with neither an antagonist or agonist bias. There is high variability in the contribution vectors at this angle. However, there appears to be no preference for any muscles in contrast to the lower angles.

#### 3.2.3 Synergy 2 : Across many angles and positions, there is a stronger bias towards the bifunctional muscles

In many individual cases, the synergy 2 pattern sets the contribution vector value of either RF or ST very high compared to the other muscles. In position 2, where the contralateral leg is bent, there is a higher average vector magnitude for RF with a lower variance than any other muscle with the exception of 0°. At 0° for both positions and at 20° in position 1 (near the extension limit) there is a strong bias to ST compared to BF. At higher angles in position 1, the ST bias appears to reduce and then vanish entirely.

#### 3.2.4 Synergy 2 : In both positions, there is a strong contribution in the antagonists at zero degrees

At 0°, in both positions, the contribution vector values for the agonist muscles are zero while the antagonist muscles show high non-zero values. This indicates that the range of activity of the antagonist muscles is reduced compared to the agonists. In position 2 at 20°, the antagonist bias is flipped to the agonists.

### 3.3 The model reproduces synergies observed experimentally

During each MIIND simulation, the cortical drive input was changed from a low to high activity to simulate the contraction behaviour. The three afferent inputs were also changed (to varying degrees depending on the desired level of afferent activity) to simulate the effect of additional load from the leg brace during the contraction. All populations produced average firing rates which were either passed to connected populations in the network or recorded for analysis. The activity of the five motor neuron populations, MN-RF, MN-VL, MN-VM, MN-ST and MN-BF, was analysed. The raw output from these populations is shown in figure 5B. The output is much smoother than the overlaid sEMG recording data due to MIIND’s simulation technique and the lack of many of the experimental sources of noise. There is undoubtedly a great deal more information available in the sEMG traces, but the model is designed only to explain how the two synergies are produced and, as we will see, the smooth rise and fall of activity is enough to do that.

The heat plots in Figure 5A show the probability density functions produced by MIIND for each population in the network. As shown in section 2.7, the density function describes the likelihood of finding a neuron from the population with a given membrane potential. The top density plot shows the state of the MN-RF population during the period before the action begins. The lower density plot shows the state when the input is maximal. In the lower density plot, there is a higher probability of finding neurons at the threshold (−51mV) indicating that the average firing rate of that population is higher. The population transitions to the top density once again after the cortical drive returns to zero. These transitions are also visible in the probability density functions of the other motor neuron populations due to the excitation from cortical drive. Therefore, for all motor neuron populations, as with the sEMG signals, the average firing rate output shows an increase to a high level of activity followed by a decrease to rest.

In the same manner as the sEMG recordings, rank 2 NMF was performed on the time series of average firing rates of the motor neuron populations in the model producing a five value muscle contribution vector and time series activation pattern for both synergies. For each trial, the maximal activity values for the afferent inputs A and C were altered and the results were compared to those of the isometric task. Figure 4 shows the results from the NMF process. For synergy 1 (Figure 4A), the activation pattern matches the shape of the descending input pattern from the cortical drive input (5 seconds of maximal activity with a 1 second ramp up and down). The five muscle contribution values are all well above zero indicating that the activation pattern is a component in the activity of all the motor neuron populations. This is in good agreement with the synergy 1 pattern observed from the sEMG data.

**Figure 4:**
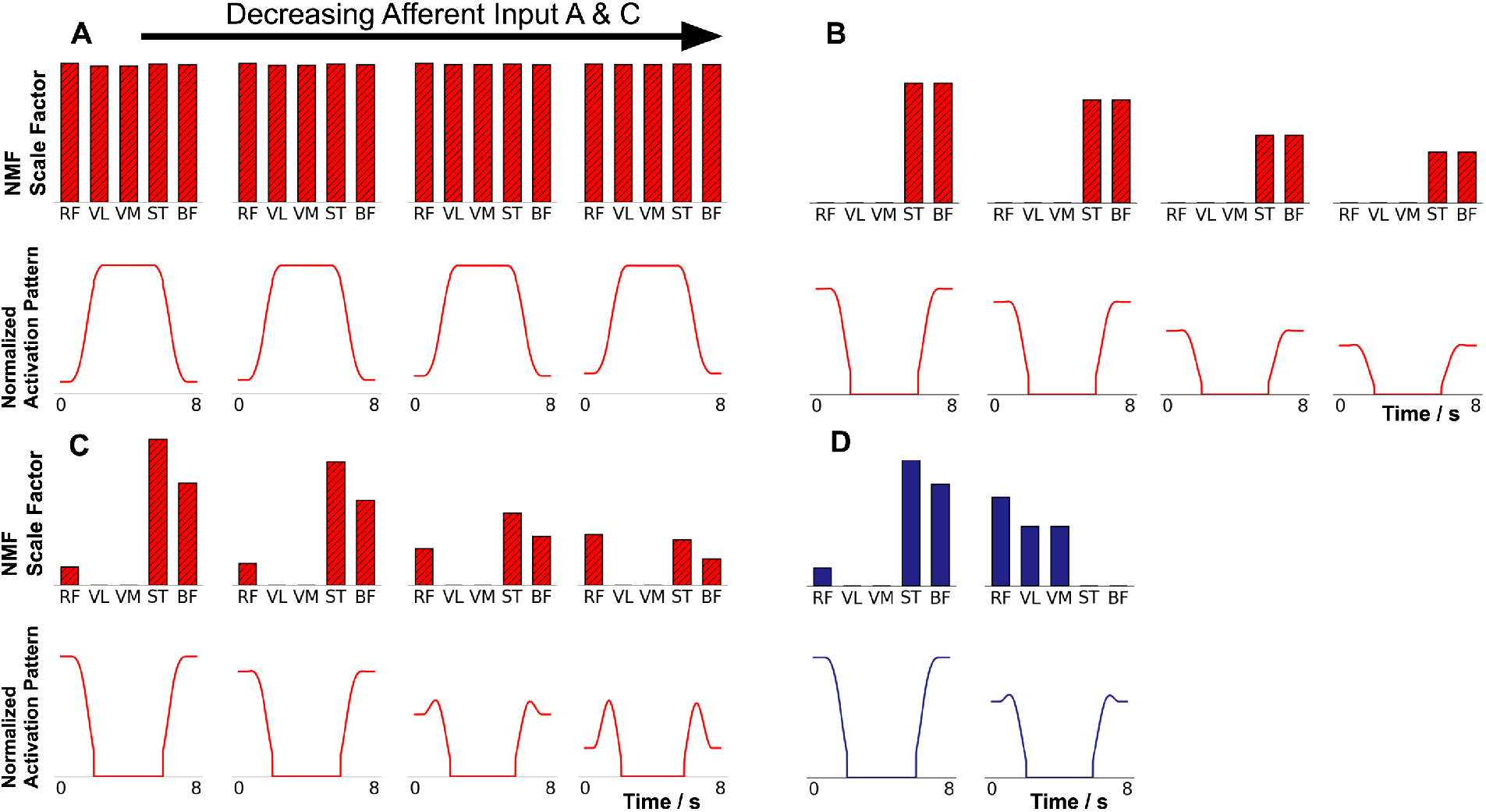
Muscle synergy features extracted using rank 2 NMF applied to the average firing rates of five motor neuron populations for different levels of input from afferent input A (1). As with the experimental results, line plots indicate the activation pattern for each synergy and bar charts indicate that pattern’s contribution to each motor neuron population’s activity. (A) Synergy 1. The activation pattern and contribution vector remains constant for all levels of afferent input.(B) Synergy 2 produced by the model in which all motor neuron populations have the same connection strength from extensor and flexor interneuron populations (MN-RF and MN-ST receive no additional bias). (C) Synergy 2 produced by the model in figure 1. The two farthest left charts show exaggerated antagonist bias due to activity from afferent input C in addition to afferent input A. (D) To reproduce the Synergy 2 patterns of position 2 at 0° (left) and 20°(right). On the right, afferent input A was further reduced to below the level of afferent input B resulting in bias towards the agonist motor neuron populations. On the left, even with a low level of afferent input A, a high afferent input C to represent maximal extension at the knee can reproduce the pattern for position 2 at 0°.

The shape of the synergy 2 time series in Figure 4B, 4C, and 4D show that, for motor neuron populations where the contribution vector values are non-zero, the resting activity of the population was higher compared to its activity during contraction. Because the raw outputs from the populations were normalised before NMF was applied, the synergy 2 time series represents both a higher resting activity and a lower maximal activity during contraction. The synergy 2 contribution values can, therefore, be considered an inverse measure of the range in activity of the original output. The similarity of the activation patterns to those from the experiment indicates that the same features are being captured and that the model provides a mechanism for producing them.

#### 3.3.1 Changing afferent inputs A or B reproduces the bias in synergy 2 between agonist and antagonist motor neuron populations

The degree to which the synergy 2 activation pattern contributes to each motor neuron population’s activity changes with the different combinations of afferent inputs. Figure 4C shows the trend in the synergy 2 contribution vector and activation pattern with decreasing afferent activity from left to right. When afferent input A is higher than B, the activation pattern contributes to the antagonist motor neuron populations significantly more than the agonists as was observed in the experimental results. The additional excitation from input A causes an imbalance in activity between the extensor interneuron population and flexor interneuron population. The resultant higher firing rate of the flexor interneuron population causes additional inhibition of MN-ST and MN-BF and excitation of MN-RF, MN-VL and MN-VM. Therefore, during the contraction, the agonist motor neuron populations have a higher maximal firing rate than the antagonist populations. This is how bias in the synergy pattern to either the agonists or antagonists is controlled. As the difference between afferent inputs A and B is reduced, the contribution of this pattern lowers until it is eliminated across all five populations.

For the sEMG recordings, at 20° in position 2, the agonist/antagonist bias is reversed. This can be reproduced in the model by increasing afferent input B above A which creates a bias towards the agonist motor neuron populations in the contribution vector as shown in Figure 4D.

#### 3.3.2 The additional connection strength to MN-RF and MN-ST reproduces the bias towards the bifunctional muscle populations in synergy 2

In position 1, the contribution vector value of ST is higher than that of BF at 0°. This bias remains at 20° then reduces at 60° before being eliminated entirely at 90°. The additional connections between the extensor interneuron population and MN-ST cause the activity of MN-ST to increase both during the rest and contraction periods. In the model, this effect can be reproduced with the additional connections between the extensor interneuron population and MN-ST. The multiplicative effect on the input to MN-ST due to the larger number of connections causes the range of activity to be reduced because the resting activity is increased more than the maximal activity. The reduced activity range is shown by a higher contribution vector value for MN-ST compared to MN-BF. This effect occurs independently of the afferent input to the flexor interneurons and so the difference is still visible even with balanced extensor and flexor interneuron populations. To eliminate the bias in line with the experimental results, the inhibitory population InhibST is required to offset the additional activity from the extensor interneuron population. This mechanism is mirrored for the MN-RF population. In position 2 except for 0°, RF has a consistently high contribution vector value. The additional connections to MN-RF from the flexor interneuron population enable a difference in the synergy pattern to that of MN-VL and MN-VM and the InhibRF population allows modulation of the bias.

#### 3.3.3 Afferent Input C provides a strong bias in synergy 2 for the antagonist muscles at zero degrees

To produce the synergy 2 patterns at 0° in both positions and 20° in position 1, all that is required is to provide a large amount of excitation through afferent input A. However, the sharp change to the synergy patterns at 60° in position 1 and 20° in position 2 indicates that the afferent input does not change linearly or there is a separate additional afferent signal causing this pattern. In both cases, this can be modelled with a separate afferent input C. In the model, at 0° in both positions, afferent input C augments the signal from input A to produce the required pattern. The additional input also allows for the synergy pattern of 0° to be achieved even if input B is higher than input A as shown in Figure 4D.

#### 3.3.4 Differences between the model synergies and experimental synergies

Figures 5C and 5D show the side by side comparison of the synergies extracted from the simulation and average synergies from the sEMG recordings for position 1. As discussed above, many of the patterns and trends are captured by the model. However, there are a few differences which we discuss here. In position 1, the model produces no contribution vector values in synergy 2 for MN-VL and MN-VM in contrast to the experimental results. The model provides no variability in the inputs to these populations and no mechanism for changing their activity in comparison to MN-RF. This is particularly obvious at 90° where, in the experimental results, there is an equal chance of seeing a high contribution vector value for all muscles which leads to the common low average vector value with high variability. The equivalent in the model is a zero vector value for all motor neuron populations in synergy 2.

**Figure 5:**
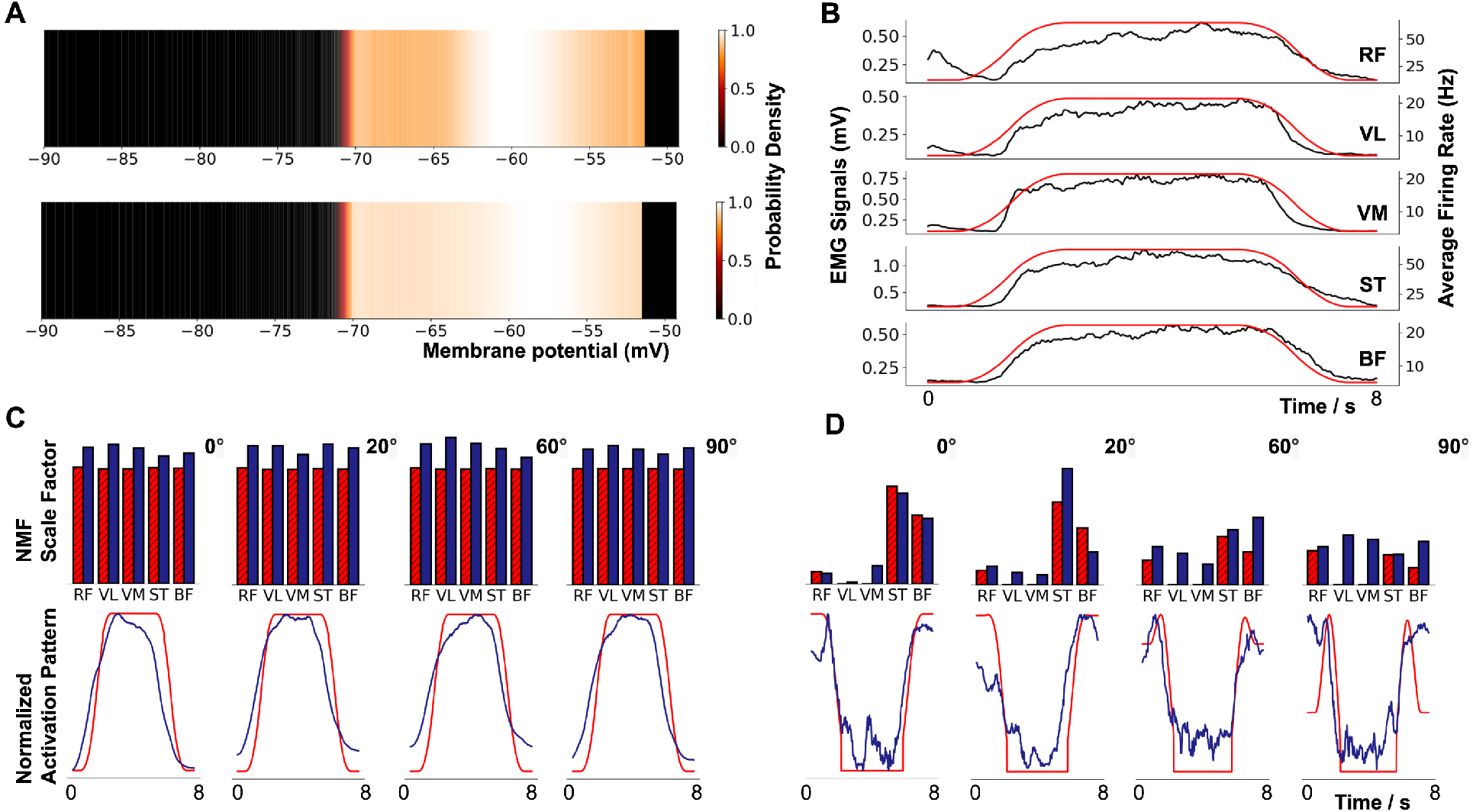
(A) The probability density function of the MN-RF population in the model before input from cortical drive (upper) and during the contraction (lower). Colour brightness indicates the probability of a neuron in the population having that membrane potential. The y axis of the plots represents an arbitrary value for simple exponential integrate and fire neurons. A higher probability at the threshold of -51mV indicates a higher average firing rate for the population.(B) The average firing rate of each motor neuron population in the model and the rectified and smoothed sEMG output for each muscle. (C and D) The muscle contribution vector and activation pattern time series for synergy 1 (C) and 2(D) for the model (hatched bars) and for experimental data from position 1 (solid bars with std errors). Indicative levels of afferent activity in the model have been used for comparison with each internal knee angle of the experiment.

The trend in agonist/antagonist synergy bias in position 1 which is perhaps extended to 20° in position 2 is not continued for 60° and 90°. The model does not currently explain the changes in synergy pattern although it is still possible to reproduce them by changing the afferent inputs and/or cortical drive to specific motor neuron populations.

## 4. Discussion

The major findings of this study show that the synergies recruited during an isometric knee extension are affected by proprioceptive feedback and that these synergies can be reproduced by a neural population model integrating afferent feedback. The synergies are well conserved across conditions and individuals, consisting of simultaneous muscle activation and the balancing of agonist-antagonist recruitment relative to internal angle of the knee. These results are further supported by findings from a novel simulation of the local spinal circuits showing proprioception from muscles contributes to synergy level organization of motor control in humans during isometric tasks. Our experimental results are largely in line with previously established literature regarding synergy recruitment in motor tasks (Saltiel et al. 2001; Tresch, Saltiel, et al. 2002; Torres-Oviedo and Ting 2007). However, even when considering only the maximal sEMG activity of each muscle at differing internal knee angles, there are conflicting results (Babault et al. 2003; Pincivero et al. 2004; Zabik and Dawson 1996). In this study, by recording from the quadriceps and hamstring muscles then using synergy analysis, we have been able to identify trends that involve agonist-antagonist interactions which are not solely dependent on the maximal sEMG output. That is, synergy 2 gives an inverse measure of the difference between the resting sEMG activity and MVC activity which is then compared across all muscles. We further identified a bias towards the bifunctional muscles, ST and RF, across the synergy 2 vector patterns. This may reflect a greater importance of these multi-joint muscles in synergistic control of movement. It could also suggest that there is more complexity to their behaviour which was not captured by NMF with rank 2.

Among studies which use synergy analysis to investigate proprioceptive effects on motor control, there is again disagreement (Roh, Rymer, and Beer 2012; Torres-Oviedo, Macpherson, and Ting 2006). We suggest that the cause of these conflicts is a level of unreliability in the NMF process itself. Barradas et al. 2020 suggest that the use of 90% as the VAF threshold for choosing the NMF rank is too lenient and important aspects of the data can therefore be missed. While 90% was used in this study, we agree that there is indeed structure in the remaining 10% which cannot be attributed to noise. However, when running NMF on this data with rank 3, we found that the activation pattern of the third synergy was visually indistinguishable from noise which gave us no indication of how it might be reproduced in the model. Furthermore, an important factor in justifying our results here is the appearance of high contribution vector values for the antagonist muscles in both synergies for rank 2. When studying synergies using NMF, it can be the case that the contribution vectors are almost entirely disjoint across synergies. When this happens, it is an indication that the NMF process has simply classified the original set of sEMGs into matching groups. Higher rank increases the likelihood of this disjoint classification occurring and is another reason why rank 2 was picked instead of rank 3. The disjoint classification problem can be confirmed if the synergy activation patterns are almost identical to the associated raw sEMGs. In our study, the high antagonist contribution values in both synergy 1 and 2, coupled with an activation pattern in synergy 2 which is starkly different to the sEMG time series, indicates that NMF has identified underlying structure in the data which we were able to use to build the model.

The mechanism of action for a given task is hard to determine in a human spinal cord due to the lack of direct recordings from these circuits, but using MIIND, we built a population network model based on biological evidence to propose the likely mechanism. The model reproduced the same synergy patterns as derived from the sEMG data. Changing the afferent inputs in the model produced the same trends in synergy contribution values as observed with increasing knee angle. This model, therefore, demonstrates how muscle synergies can be encoded in neural population circuits and that synergy analysis of experimental data can be used to directly drive model development.

In simulations, instead of using the traditional technique of direct simulation of individual neurons, we have demonstrated the use of the MIIND simulation package, a software environment allowing easy simulation of populations of neurons. MIIND requires only the definition of connectivity at the population level, making it easy to set up and adjust a population network during development. Parameter tweaking is an inevitable part of the modelling process requiring cycles of adjustment followed by simulation. Reducing the need for adjustments to the neuron model itself was one reason why we used the simple exponential integrate and fire instead of a more complex Hodgkin-Huxley style neuron. We were able to reproduce the desired synergy patterns without the need for such complexity. While building the network model we experimented with different connection configurations between populations. MIIND’s XML style code, used to describe the network, made it simple to add, remove or adjust connections, as well as to add further populations for the RF and ST bias. For one dimensional neuron models, MIIND can simulate a population network with much greater speed than direct methods and this allowed simulations to be run on a local machine without the need for high-performance computing, significantly improving the turnaround time between changing and testing the model. From our experience here, we advocate the use of simple neuron models where appropriate, i.e. reduce the dimensionality of the neural model as far as possible. First, this increases simulation speed and second, this forces thinking about which are the essential neuronal mechanisms before simulation starts.

This model’s similarity to previously studied spinal circuits (Hultborn et al. 1987; Shevtsova et al. 2016; Pierrot-Deseilligny and Burke 2005), supports conclusions (Saltiel et al. 2001; Dominici et al. 2011) that synergy encoding takes place in the spinal cord. Sohn and Ting 2016 describe a cat musculoskeletal model from which synergies can be derived at different postures. Their model eliminates the possibility that synergies derive from the biomechanical constraints of the system alone but suggest a combination of factors which could include neural control mechanisms as we have demonstrated here. Inspection of the synergies extracted during NMF analysis matches well with expected muscle interactions, with synergy 1 reflecting coordinated contraction of all muscle groups and synergy 2 reflecting the agonist-antagonist pairing of the hamstrings and quadriceps. Supraspinal input was provided equally to the motor neuron populations (with the exception of RF) and extensor and flexor interneuron populations so specific bias, e.g. towards the antagonist muscles, was only introduced through the circuitry of the model itself leading to the patterns observed in synergy 2. During this study, it became clear that altering the circuitry of the model (and the level of afferent input) affected the amplitude of activity of each motor neuron population at rest and during maximal contraction. The contribution value of synergy 2 in the simulation was found to be inversely proportional to the difference between the maximal and minimal activity which can also be observed in the experimental results.

The similarity of each synergy’s activation pattern across the experimental and simulated results indicates that the same synergies are being recruited at each angle. The changes observed in synergy 2 at different internal angles of the knee demonstrates a clear effect of proprioception on synergy contribution to each muscle. In position 2, with the contralateral hip flexed, the trends with internal knee angle are less apparent. In position 2, passive insufficiency is reduced which may indicate that the afferent signals identified here are most active when the muscles are under higher tension. If we ignore the special case of 0 degrees in position 2, the synergy 2 pattern at 20° (bias towards the agonist muscles) could be a continuation of the trend observed in position 1. The model is still able to reproduce the synergies observed at each angle. It can be said that the quadriceps and hamstring activity in synergy 2 becomes balanced at a lower angle (20° instead of 60°) in position 2. This further reinforces that the changes in synergy recruitment reflect the influence of afferent feedback due to change in relative muscle stretch.

Inputs A and B of the model were used to control the general balance of interneuron activity between the flexor and extensor motor neuron populations in order to produce the results between 0°and 90°in both positions. Afferent input C was used to provide additional excitation to the flexor interneuron population for the strong agonist bias in the synergy 2 pattern at 0° for both positions (and 20° in position 1). At 0°, there is a mechanical restriction on further knee extension and it is reasonable to assume that a neural restriction might also be in place. However, this would imply a reduced activity of extensor motor neurons and increased activity of flexor motor neurons which was not observed or produced in the model. Babault et al. 2003 showed a similar trend in surface sEMG activity with changing knee angle and hypothesised that a greater activity near to maximal extension could be caused by a neural mechanism to overcome the mechanical restriction rather than to avoid it. It is likely that additional neural circuitry exists between afferent input C and the flexor interneuron population which integrates the afferent signals and cortical control signals to produce a form of override.

One way to evaluate the success of a model is to consider how it might be integrated into larger models to answer different research questions. Central pattern generator (CPG) models are constructed from mutually inhibiting populations of bursting neurons to produce an oscillating pattern of activity for driving rhythmic behaviours in many species and areas of the body. The lowest layer of the three layer CPG model for driving fictive locomotion in cats (McCrea and Rybak 2008) has many similarities to the model proposed here. Both include mutually inhibiting populations of interneurons, with a proprioceptive input. The use of separate inhibitory populations for controlling bifunctional muscles was also first demonstrated by Shevtsova et al. 2016. Integrating our model would require a decision about whether the cortical drive should be mediated by the higher layers of the CPG or if it should bypass them. Answering this question would give insight into how voluntary movements and cycling (CPG controlled) movements are performed by the same set of neural circuits.

In a more clinical area of research, multiple studies have shown that Parkinson’s disease affects proprioception, reducing the ability to accurately sense limb position (Ribeiro Artigas et al. 2016; Mongeon, Blanchet, and Messier 2009; Adamovich et al. 2001). While there appears to be little effect from Parkinson’s disease on the muscle spindles or afferent pathways at the level of the spinal cord, the source of the effect in the brain remains unknown. A candidate area in the brain is the Supplementary Motor Area (Jacobs and Horak 2006). By combining a neural model of this area with the spinal model proposed here, a better prediction about the influence of Parkinson’s Disease on proprioception could be made in the future.

In conclusion, despite a level of disagreement in the literature about the effect of proprioception on muscle activity in isometric tasks and about the effect of proprioception on synergies derived from muscle activity, the synergy analysis from our experiment clearly shows that there is an effect. In an isometric knee extension task, as the internal knee angle approaches 90°, the activity of the agonist and antagonist muscles becomes more balanced. Near the limit of extension, our model predicts an additional proprioceptive signal which adds a further bias to the antagonist muscles. We caution that synergy analysis should be used with an understanding that the process can give conflicting results from the same data set but indicators such as overlapping contribution vectors and markedly different activation patterns between synergies can help to support conclusions. By building a computational model of neural circuitry which can produce the same synergies as observed experimentally, we have also shown how synergy analysis can be used to help develop models of underlying neural mechanisms for motor tasks. We recommend the use of simple neuron models such as exponential integrate and fire where ion channel dynamics are not required to explain observations. Finally, the model we have demonstrated here should be integrated into larger models of motor control to add the observed influence of proprioception at the spinal cord level.

